# Genetic accessibility, not superior fitness, determines which anti-HBs antibody escape variants are selected in patients

**DOI:** 10.64898/2026.10.06.756817

**Authors:** Yingpu Yu, Eleftherios Michailidis, Leon L Seifert, Mengyin Zhang, Chenhui Zou, Catherine A Freije, Georgios Dangas, Antonis Athanasiadis, Andreas Puschnik, Audrey Hanson, Evelyn Rocha, Yi-Pei Chen, Xupeng Hong, Corrine Quirk, Yichen Zhou, Michael A Schmid, Florian A Lempp, Charles M Rice, William M Schneider, Ype P de Jong

## Abstract

**Background and Aims:** Genetic diversity within the hepatitis B virus (HBV) surface antigen (HBsAg) drives the emergence of immune escape variants, but studying this diversity *in vivo* has been limited by the difficulty of establishing infection from patient isolates. We aimed to perform unbiased *in vivo* screening of variants in the HBs antigenic loop for antibody escape and fitness, using synthetic DNA launch of HBV in human hepatocyte chimeric mice.

**Approach and Results:** Direct intrahepatic injection (IHI) of barcoded, recombinant cccDNA established infection from up to 32,000 independent launch events of a genotype A strain and successfully launched clinical consensus sequences across the major genotypes. Using IHI, we performed deep mutational scanning of 4,608 codon variants across the HBs antigenic loop. Under control antibody treatment, diverse variants expanded, revealing differences in variant fitness in the absence of adaptive immune pressure. Anti-HBs monoclonal antibody treatment suppressed wild-type viremia while permitting breakthrough of resistant variants from the library. Sequencing identified strong enrichment for substitutions at G145 (G145Q, G145H, G145R). Notably, G145Q, rather than the clinically dominant G145R, showed the greatest fitness advantage, indicating that G145R’s clinical prevalence does not reflect superior intrinsic fitness.

**Conclusions:** Unbiased *in vivo* screening reveals that clinically observed HBV escape mutations reflect which substitutions are genetically accessible by single nucleotide change and compatible with the overlapping polymerase reading frame, rather than which confer optimal fitness. This approach establishes a platform for high-resolution mapping of viral variant fitness under antiviral selection pressure.

## Introduction

Hepatitis B virus (HBV) is a major human pathogen causing approximately 649,000 annual deaths worldwide(1). Although effective prophylactic vaccines are available, therapeutic options for chronically infected individuals remain limited. Nucleot(s)ide analogs suppress viral replication across genotypes and variants but rarely achieve a functional cure, typically requiring lifelong treatment. Interferon-α-based therapies can induce a functional cure in a substantial proportion of HBeAg-positive individuals infected with HBV genotype A (HBV-A), but they are less effective in other genotypes or in HBeAg-negative patients (2–6). Several new classes of antivirals are under clinical development, including neutralizing antibodies (nAbs) targeting the surface antigen (HBsAg). Because nAb efficacy will depend on how easily HBV can escape, new methods are needed to interrogate viral sequence diversity *in vivo* and anticipate resistance.

A central barrier to understanding HBsAg escape has been the narrow repertoire of viral variants accessible to preclinical study. HBV has restricted tropism for human and chimpanzee hepatocytes, and historically, cell culture studies have relied on plasmid DNA transfection systems and a limited number of human hepatoma cell lines containing integrated HBV DNA (7, 8). Following identification of the viral entry receptor, the sodium taurocholate co-transporting polypeptide (NTCP), human hepatoma cells engineered to overexpress NTCP have increasingly been used to study infection by clinical HBV isolates (9–11) but these systems capture only the specific variants present in the infecting isolate rather than the broader mutational space relevant to antibody escape. *In vivo*, the discontinuation of chimpanzee studies has shifted most investigations toward mice (12), yet because of HBV’s narrow tropism, rodents cannot support the complete viral life cycle. Germline HBV-transgenic mice or transient HBV expression via plasmid DNA delivered using hydrodynamic tail vein injection (HDI) or adeno-associated virus vectors are commonly used (13, 14), but these approaches similarly rely on predefined, often single, sequences and do not permit interrogation of the full HBV life cycle in the context of a genetically diverse population. Hepatocyte chimeric mouse models (15) generated by xenografting primary human hepatocytes into immunodeficient mice permit authentic HBV infection and completion of the full viral life cycle (16), offering the closest *in vivo* approximation of natural infection. However, these models exhibit variable infectivity, and for unknown reasons, not all patient-derived HBV isolates can establish infection, even when hepatocyte engraftment is high. Therefore, HBV-positive serum is serially passaged between chimeric mice, further narrowing the isolates available for study. Defining which variants can emerge under antiviral pressure requires a genetically diverse input population and a system with sufficient infection efficiency to interrogate this diversity. While diverse virus populations are readily obtained from patients, existing models establish infection too inefficiently to sample them. As a result, no existing system has permitted unbiased, quantitative interrogation of HBV variants that can emerge under antiviral pressure, a critical gap for anticipating resistance to compounds now in clinical development.

Here, we sought to comprehensively map all single-codon variants in the HBs antigenic loop that can escape neutralization by a potent anti-HBs nAb. To accomplish this, we first developed multiple DNA launch platforms in human hepatocyte chimeric *Fah^−/−^* NOD *Rag1^−/−^ Il2rg^null^* (huFNRG) mice (17) and compared their efficiencies using synonymous mutations as sequence barcodes that maintain amino acid identity while enabling quantitative detection by sequencing. Among these, recombinant covalently closed circular (rccc)DNA delivered by direct intrahepatic injection (IHI) was the most efficient, enabling the simultaneous launch of thousands of viral genomes. We then leveraged this approach to perform *in vivo* deep mutational scanning (DMS) of the hepatitis B surface (HBs) antigenic loop (AGL), allowing us to define viral sequence constraints *in vivo* under a nonselective condition using a control monoclonal neutralizing antibody (nAb), as well as to identify escape variants in the presence of an HBs nAb.

## Results

### Establishing HBV infection in human hepatocyte chimeric mice

A key challenge in studying HBV variant biology is reliably establishing infection in human hepatocyte chimeric mice with diverse viral isolates. Building on prior work (18–20) in which we successfully infected huFNRG mice with an HBeAg^−^ patient-derived HBV-C isolate (HBV-C_Y) that retained high specific infectivity upon passage (**Sup Fig S1a**), we attempted to expand our repertoire of infectious strains. However, conventional approaches, including intravenous challenge with patient plasma and infection with hepatoma cell line supernatants under various conditions, failed to achieve viremia (**Sup Fig S1b–d**). Only very high inocula of concentrated supernatant from HepDE19 hepatoma cells (HBV-DE19) were able to infect mice (**Sup Fig 1e**). These results indicated that patient-derived plasma and cell culture-derived virus often lack sufficient specific infectivity to reliably infect huFNRG mice, possibly due to inhibitory factors such as antibodies or excess HBsAg in serum, and/or inherently low specific infectivity.

**Sup Figure S1:**
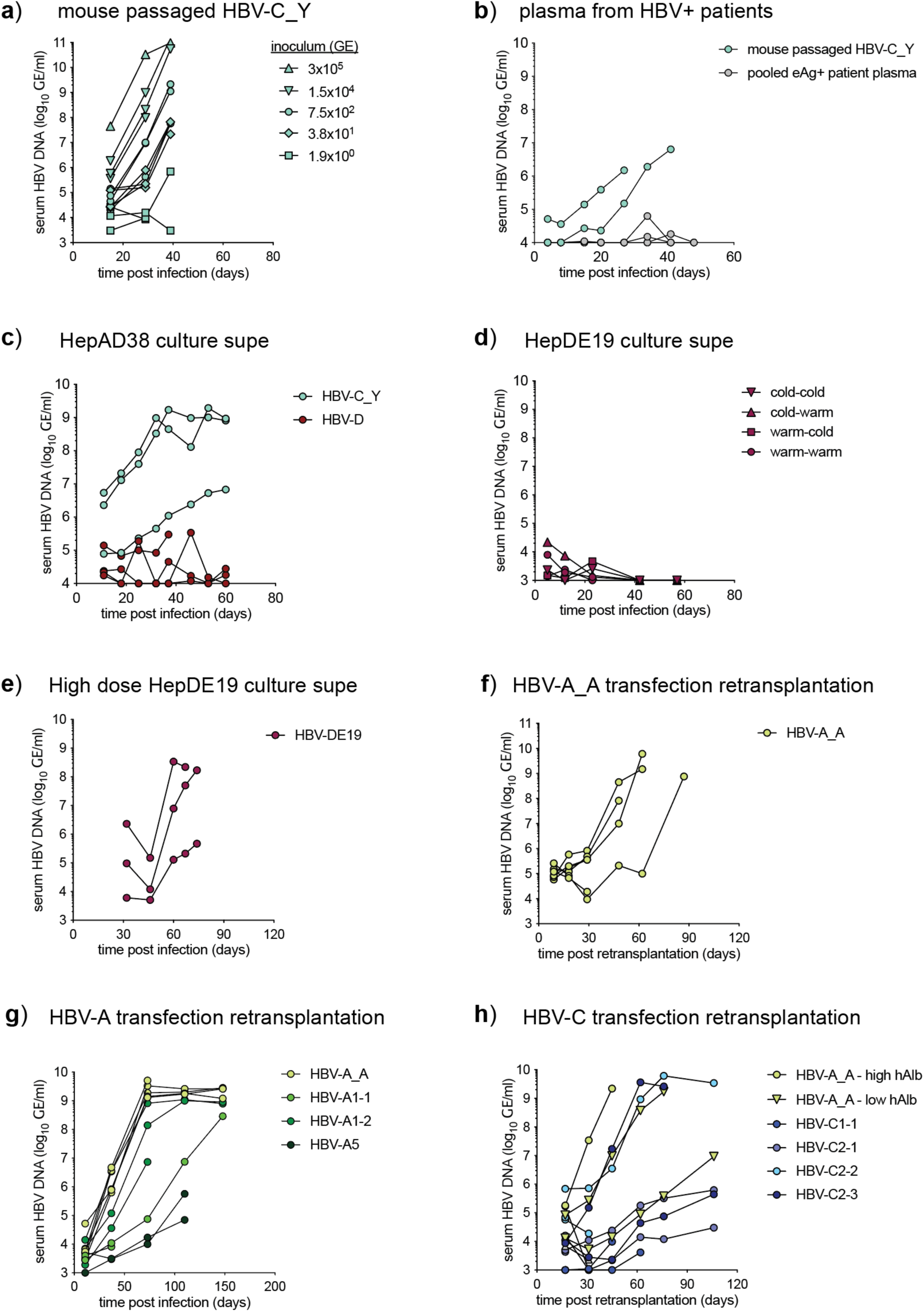
Launching HBV isolates in huFNRG mice. **a)** huFNRG mice were challenged intravenously with HBV-C_Y containing serum at 20-fold dilutions spanning an inoculum range from 1.9×10^0^ to 3×10^5^ genome equivalent (GE) per mouse, and HBV DNA in mouse serum was quantified over time. All mice in the 3.8×10^1^ GE group and 1 of 3 mice in the 1.9×10^0^ GE group became viremic. **b)** huFNRG mice were challenged intravenously with pooled plasma from four HBeAg^+^ blood donors at >5×10^8^ GE per donor, or mouse-passaged HBV-C_Y serum. None of these four patient isolates established viremia over this time course. **c)** huFNRG mice were challenged intravenously with cell culture supernatant from HepAD38 cells containing 4×10^6^ or 2×10^8^ GE, or mouse-passaged HBV-C_Y serum. HepAD38 supernatants did not induce viremia over this time course, regardless of dose. **d)** Culture supernatants from HepDE19 cells were stored at 37°C or at 4°C prior to injection of 1×10^8^ GE into huFNRG mice. This did not result in viremia over this time course. **e)** Culture supernatants from HepDE19 cells were concentrated using polyethylene glycol prior to injection of 3×10^9^ GE per huFNRG mouse. After an initial decline, all mice became viremic. **f)** mpPHH cultures were transfected with HBV-A_A rcccDNA prior to retransplantation into huFNRG mice. Four of five mice became viremic. **g**-**h**) The transfection and retransplantation protocol was used to launch different consensus sequences from HBV-A and HBV-C subgenotypes, showing variable success between and within isolates (high hAlb = 10 mg/ml, low hAlb = 2 mg/ml). For **a**-**h**) symbols represent individual mice.

**Figure 1:**
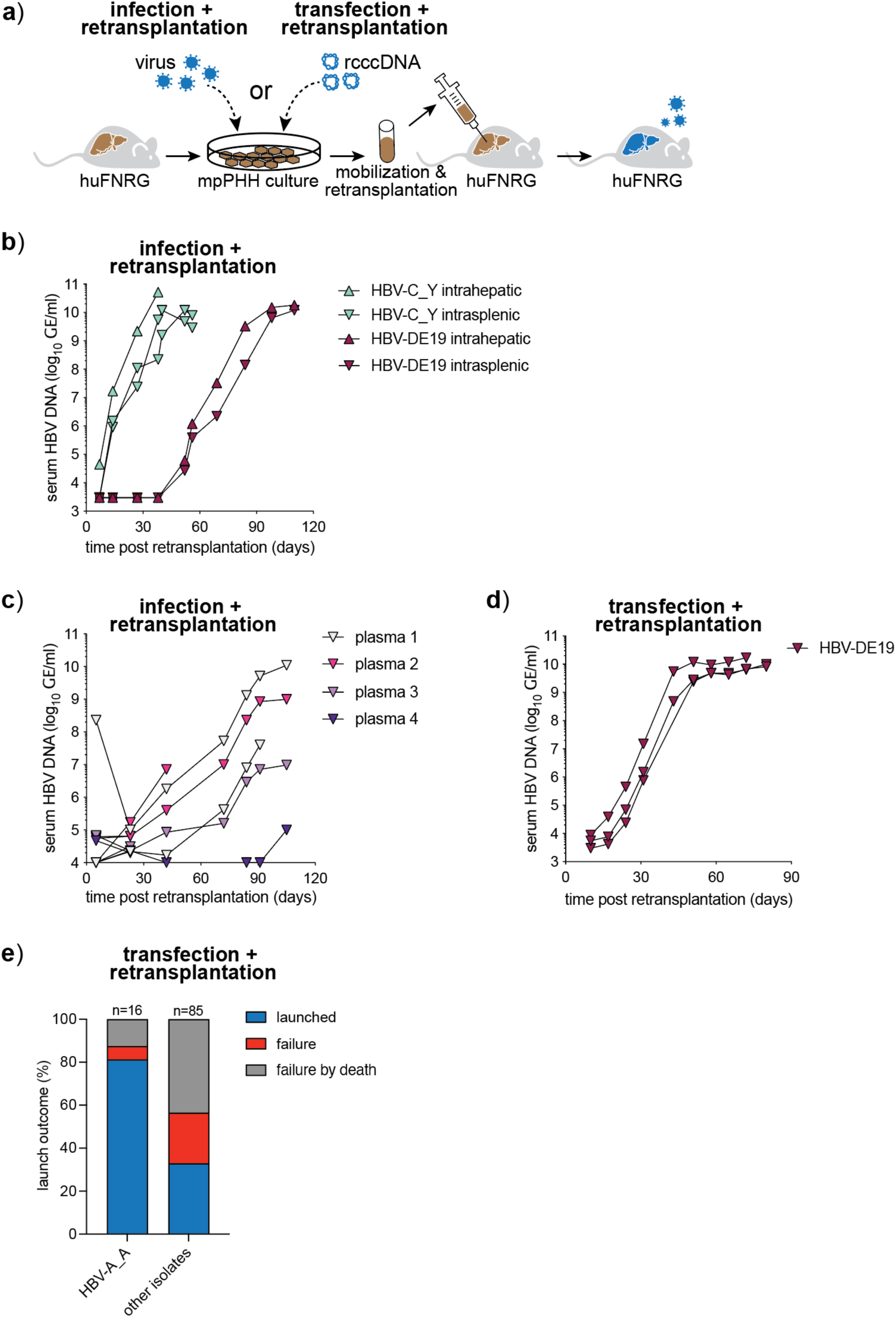
Launching HBV strains using hepatocyte re-transplantation. **a)** Experimental schematic of the hepatocyte re-transplantation protocols. Mouse-passaged primary human hepatocytes (mpPHH) were isolated from huFNRG mice, plated in 2-dimensional cultures, and either infected with HBV virus or transfected with rcccDNA prior to mobilization and transplantation in already humanized huFNRG mice. **b)** mpPHH cultures were infected with huFNRG serum containing HBV-C_Y or HepDE19 cell culture supernatants, mobilized, and re-transplanted into huFNRG mice via intrahepatic or intrasplenic delivery. HBV DNA in mouse serum was quantified over time. **c)** mpPHH cultures were infected with HBeAg+ patient plasma prior to retransplantation into huFNRG mice. HBV DNA in mouse serum was quantified over time (symbols indicate individual mice). **d)** mpPHH cultures were transfected with HBV-DE19 rcccDNA prior to retransplantation into huFNRG mice. HBV DNA in mouse serum was quantified over time. **e)** Cumulative launch outcomes of rcccDNA transfection and retransplantation across 17 isolates (“launched” = 2 sequentially rising HBV DNA viremia measurements above the limit of quantification, “failure” = absence of two sequentially rising viremia measurements by 12 weeks after re-transplantation, and “failure by death” = mice died before two sequentially rising viremia measurements). For **b**-**e**) symbols are individual mice.

To overcome this barrier, we developed two retransplantation-based strategies using mouse-passaged primary human hepatocytes (mpPHH) (21) as carriers (**Fig 1a**). In the first, termed infection and retransplantation (IRT), mpPHH cultures were infected with HBV *in vitro*, then mobilized and transplanted into huFNRG mice. This approach successfully established viremia with HBV-DE19 regardless of injection route (**Fig 1b**), and when applied to the four HBeAg^+^ clinical isolates that had previously failed to infect huFNRG mice by intravenous challenge, three of four caused viremia within 90 days (**Fig 1c**). In the second strategy, named transfection and retransplantation (TRT), we replaced the infection step with transfection of recombinant covalently closed circular DNA (rcccDNA), generated by enzymatic digestion and ligation of HBV plasmids. This approach offers critical advantages: it enables the generation of infectious virus from consensus sequences and allows for the engineering of isogenic variants. TRT successfully initiated infection with HBV-DE19 (**Fig 1d**) and an HBV-A strain (22) (HBV-A_A, **Sup Fig S1f**), which was infectious in huFNRG mice when delivered as pregenomic RNA using a similar approach (23). To determine the generalizability of this approach, a panel of 25 consensus sequences (24) spanning HBV genotypes A–F was synthesized as DNA and launched using TRT, using HBV-A_A as a positive control. This initiated infection in 28 of 85 mice (33%), with HBV-A_A achieving the highest success rate (13 of 16 mice, 81%) (**Fig 1e**; **Sup Fig S1g–h**; **Sup Table 2**). While these retransplantation methods substantially expand the range of HBV isolates amenable to study in huFNRG mice, and TRT allows efficient launch from DNA for some isolates, infection efficiency varied across strains, highlighting remaining challenges for general applicability.

### Silent mutants to quantify DNA launch efficiency

The variable success of the TRT method prompted us to explore two alternative strategies for launching HBV infection from DNA in huFNRG mice: direct intrahepatic injection (IHI) of rcccDNA and intravenous transfer of serum from non-humanized NRG mice that had undergone hydrodynamic tail vein injection (HDI) with HBV plasmid DNA, a commonly employed method for inducing viremia in immunocompetent mice (25) (**Fig 2a**). IHI of rcccDNA into highly humanized huFNRG mice (∼90% chimerism) produced rapid and consistent viremia with HBV-A_A and HBV-D3, the latter of which had caused viremia in only 1 of 3 TRT mice (**Sup Table 2**), while a catalytically dead polymerase mutant (YMHD) mouse did not become viremic, as expected (**Sup Fig S2a**). Similarly, serum harvested from NRG mice 21 days after HDI with HBV-A_A 1.3× plasmid DNA (after residual non-virion DNA contamination had cleared as determined by YMHD controls; **Sup Fig S2b**), successfully infected 4 of 5 huFNRG mice (**Sup Fig S2c**). Both approaches thus proved effective at launching HBV infection from DNA, with early and consistent viremia kinetics after IHI suggesting a high number of initial launch events.

**Figure 2:**
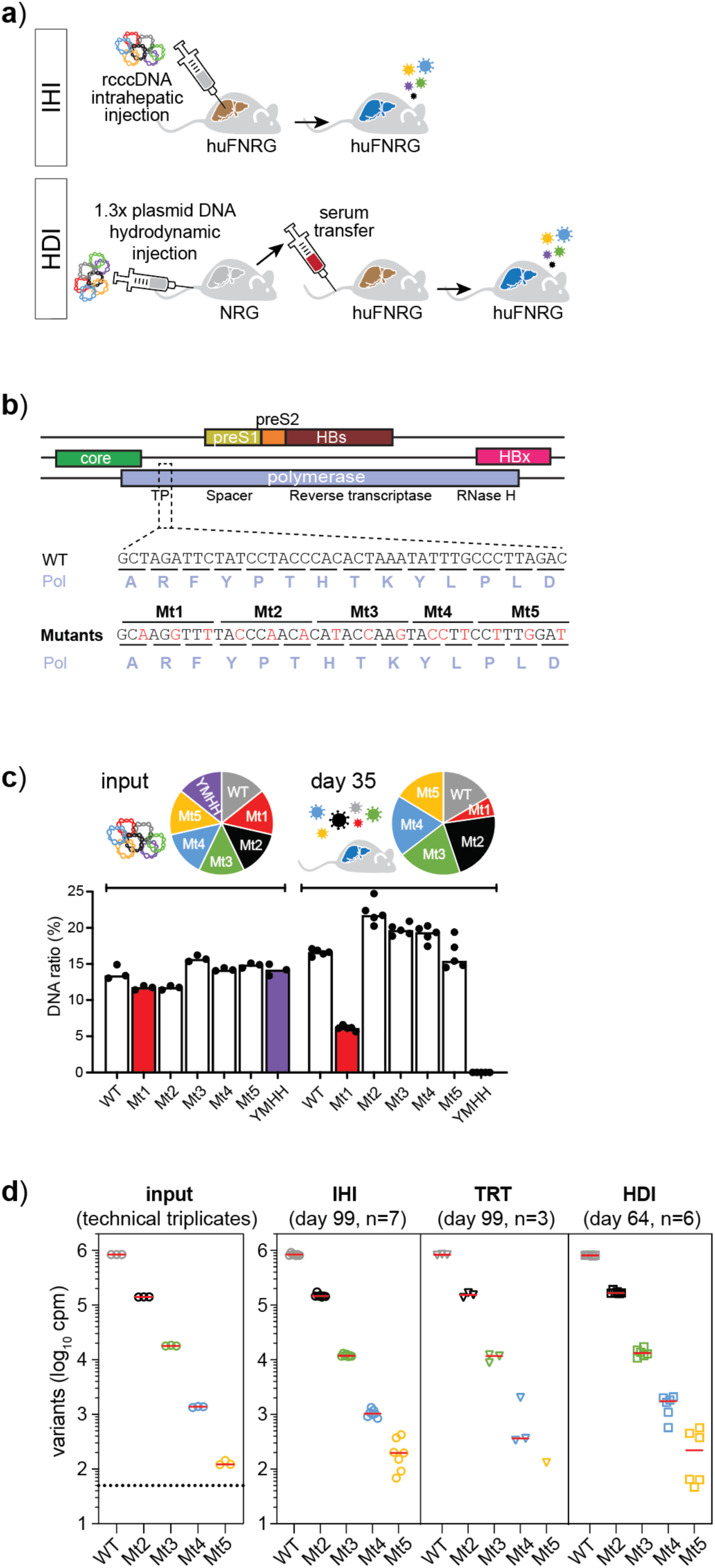
Silent mutants to quantify DNA launch efficiencies. **a)** Schematic of the recombinant covalently closed (rccc)DNA intrahepatic injection (IHI) and 1.3x plasmid DNA hydrodynamic tail vein injection (HDI) protocols. **b)** HBV genome organization with overlapping open reading frames in color. Mutants (Mt) 1-5 were created in a region of the TP domain without overlapping reading frames. Mt 1-5’s synonymous nucleotide changes (red) preserve the polymerase amino acid sequence (grey). **c)** Quantification of input rcccDNA and HBV DNA isolated from huFNRG serum at day 35 was measured by amplicon sequencing (input symbols are technical triplicates, day 35 symbols are individual mice). **d)** Quantification of mutants in the input rcccDNA and HBV DNA isolated from huFNRG mice that underwent IHI, transfection and retransplantation (TRT), and HDI (input symbols are technical triplicates, symbols for IHI, TRT, and HDI are individual mice).

**Sup Figure S2:**
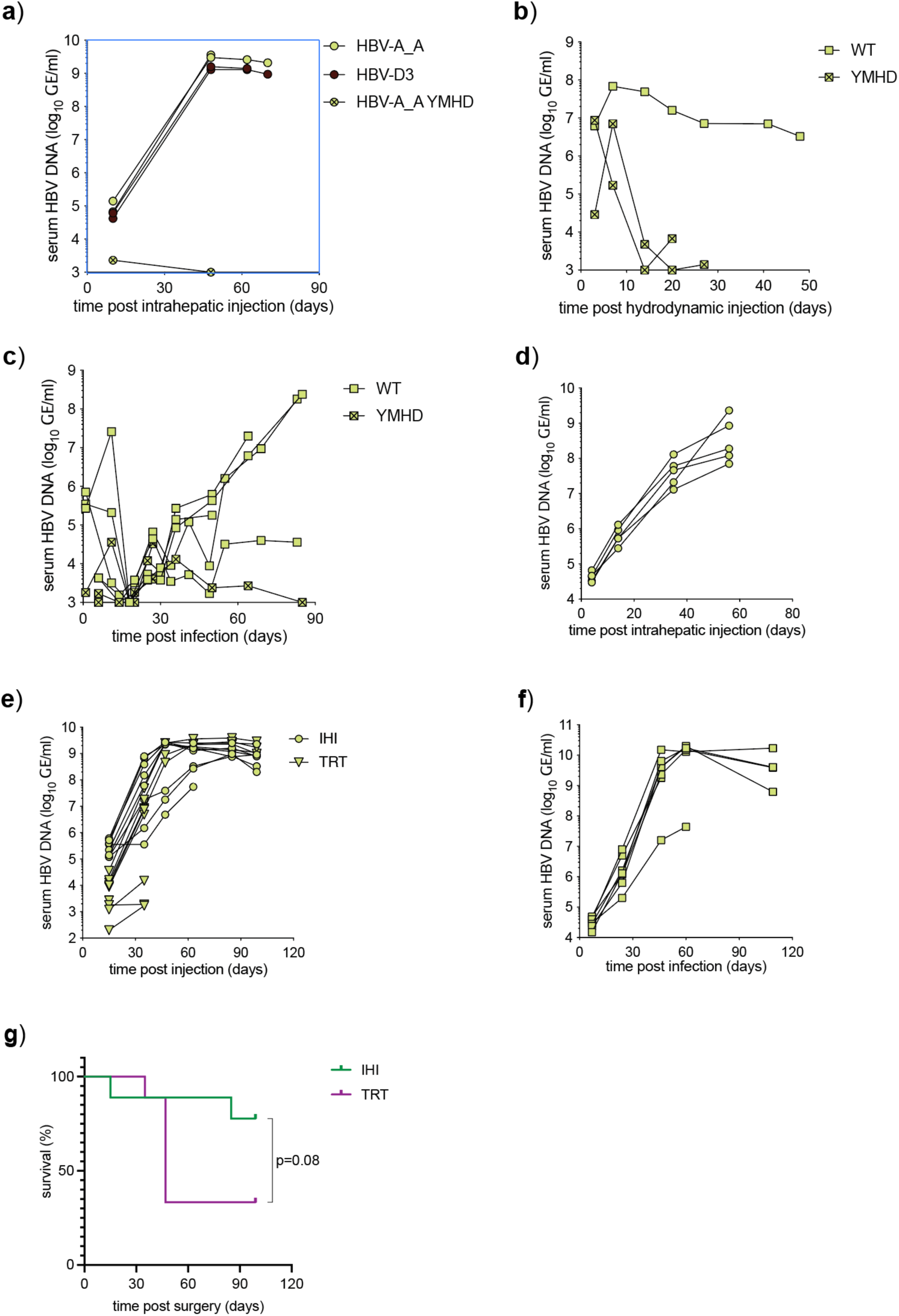
Silent mutants to quantify launch methods. **a)** huFNRG mice underwent IHI with HBV-A_A, HBV-D3 or a polymerase dead (YMHD) HBV-A_A isolate. HBV DNA in mouse serum was quantified over time (symbols represent individual mice). **b)** NRG mice underwent hydrodynamic tail vein injection with WT or catalytically dead polymerase (YMHD) 1.3x HBV plasmid DNA and HBV DNA was quantified over time (symbols represent individual mice). **c)** huFNRG mice were challenged with serum from NRG mice 21 days after hydrodynamic tail vein injection containing 7×10^5^ GE/mouse (symbols represent individual mice). **d)** An equal ratio of HBV-A_A WT, Mt1-5, and polymerase dead (YMHH) control rcccDNA was launched in huFNRG mice through IHI. HBV DNA in mouse serum was quantified over time (symbols represent individual mice). **e)** HBV DNA in mouse serum was quantified over time following launch using the following mixture: Mt2-5 were diluted 10-fold into WT rcccDNA, starting with Mt2 at 1:10 and ending with Mt5 at 1:10,000. This was transfected into mpPHH prior to retransplantation (TRT) or delivered through IHI (symbols represent individual mice). **f)** huFNRG mice were infected with serum from NRG mice that underwent HDI with 1.3x plasmid DNA containing Mt2-5 at a 10-fold dilution similar to **e**). HBV DNA in mouse serum was quantified over time (symbols represent individual mice). **g)** Groups of nine huFNRG mice underwent intrahepatic rcccDNA injection (IHI) or mpPHH transfection and retransplantation (TRT) and were monitored for survival (Mantel-Cox test).

To quantify launch efficiency across methods, we engineered five synonymous “barcode” mutants (Mt1–5) of HBV-A_A, each carrying three nucleotide substitutions in a region of the polymerase TP domain that does not overlap with other open reading frames (ORFs), providing unambiguous identification by sequencing while minimizing the risk of altering viral fitness (**Fig 2b**). To verify phenotypic neutrality, equal-ratio IHI of Mt1–5, WT, and a polymerase-dead control (YMHH) were injected into huFNRG mice (**Sup Fig 2d**), and deep sequencing of viremic serum DNA confirmed that Mt2–5 were phenotypically silent with ratios remaining close to WT. By contrast, Mt1 declined, indicating a fitness cost (**Fig 2c**). We excluded Mt1 and proceeded to spike in Mt2–5 at successive 10-fold dilutions into WT rcccDNA (i.e., Mt2 at 1:10 through Mt5 at 1:10,000) and compared detection after IHI, TRT (**Sup Fig S2e**), and HDI serum transfer (**Sup Fig S2f**). After IHI, the rarest variant (Mt5, spiked at 1:10,000) was detected in 7 of 7 mice by day 99 (**Fig 2d**). HDI serum transfer showed comparable efficiency, with Mt5 detectable in all surviving 6 mice at day 64. By contrast, Mt5 was detected in only 1 of 3 surviving TRT mice at day 99 (**Fig 2d**) while the remaining mice did not survive (**Sup Fig 2g**).

These results demonstrate that synonymous barcode mutants enable quantitative comparison of DNA launch methods, and establish that both rcccDNA IHI and passaged HDI serum launch HBV-A_A in huFNRG mice more efficiently than TRT. Because IHI and HDI also appeared to be better tolerated, we adopted these approaches for subsequent experiments.

### Identifying the method that can launch most HBV variants from DNA

Having established that silent mutant barcodes can quantify DNA launch efficiency, we next used these to define which approach yielded the most DNA-derived launch events: rcccDNA IHI, 1.3× plasmid DNA IHI, and 1.3× plasmid DNA HDI (**Fig 3a**). If 1.3× plasmid DNA proved to be as efficient as rcccDNA, it would eliminate the labor-intensive step of generating rcccDNA. To enable a rigorous comparison, we first needed to expand the barcode panel and optimize detection sensitivity for rare variants.

**Figure 3:**
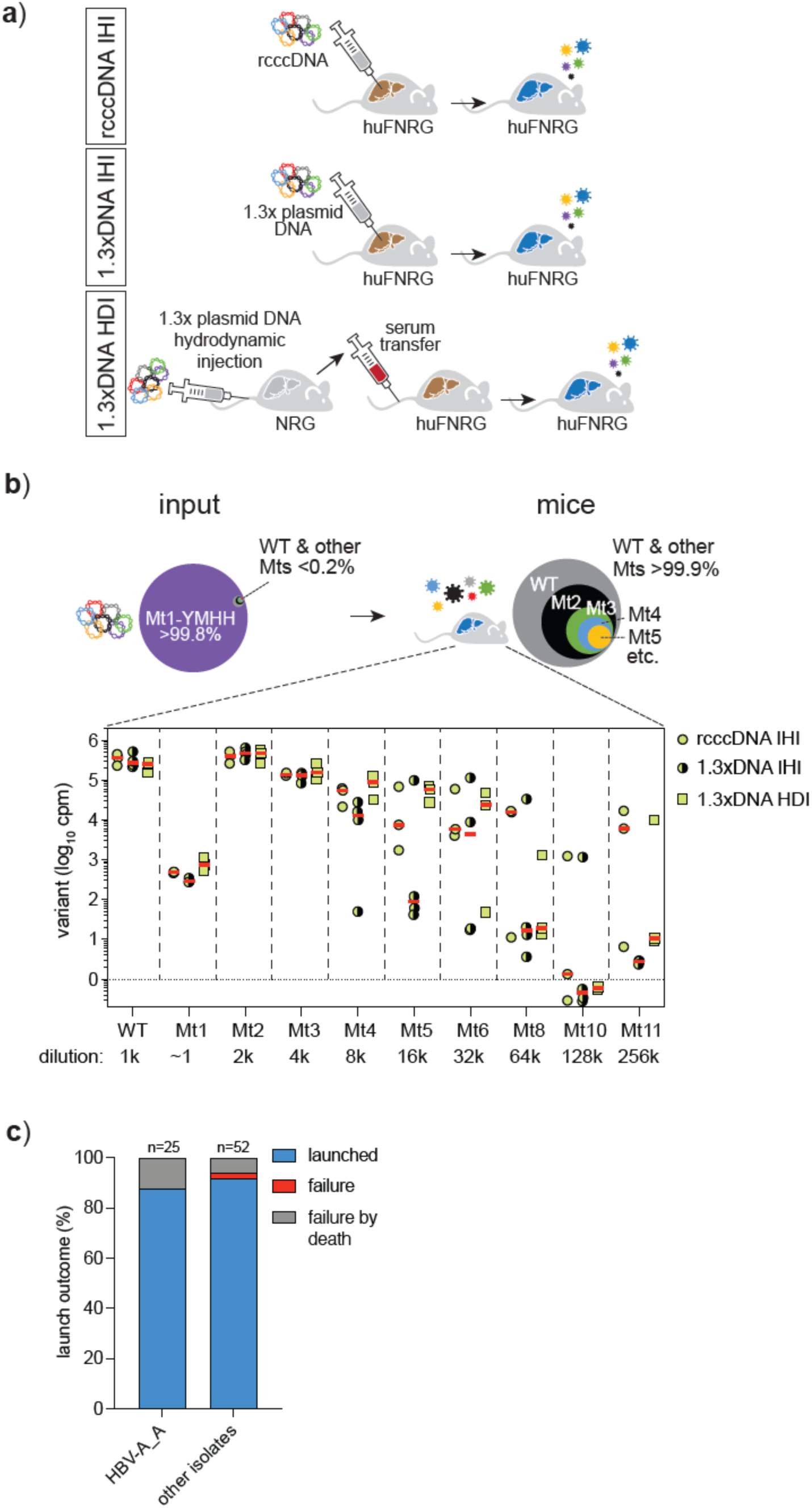
rcccDNA intrahepatic injection is optimal for launching low-frequency variants. **a)** Schematic of three methods that were further tested to quantify DNA launch efficiency. In addition to the previously established recombinant covalently closed (rccc)DNA intrahepatic injection (IHI) and 1.3x plasmid DNA hydrodynamic tail vein injection (HDI) protocols, we tested 1.3x plasmid DNA IHI. **b)** Amplicon sequencing of mutants in mice at peak viremia showed a gradual decrease in detected mutants (cpm = count per million; symbols: individual mice). **c)** rcccDNA IHI was used to launch infection of 14 isolates across genotypes B-F, with a cumulative success rate that was comparable to HBV-A_A (“launched” = 2 sequentially rising HBV DNA viremia measurements above the limit of quantification, “failure” = absence of two sequentially rising viremia measurements by 12 weeks after IHI, and “failure by death” = mice died before two sequentially rising viremia measurements)

**Sup Figure S3:**
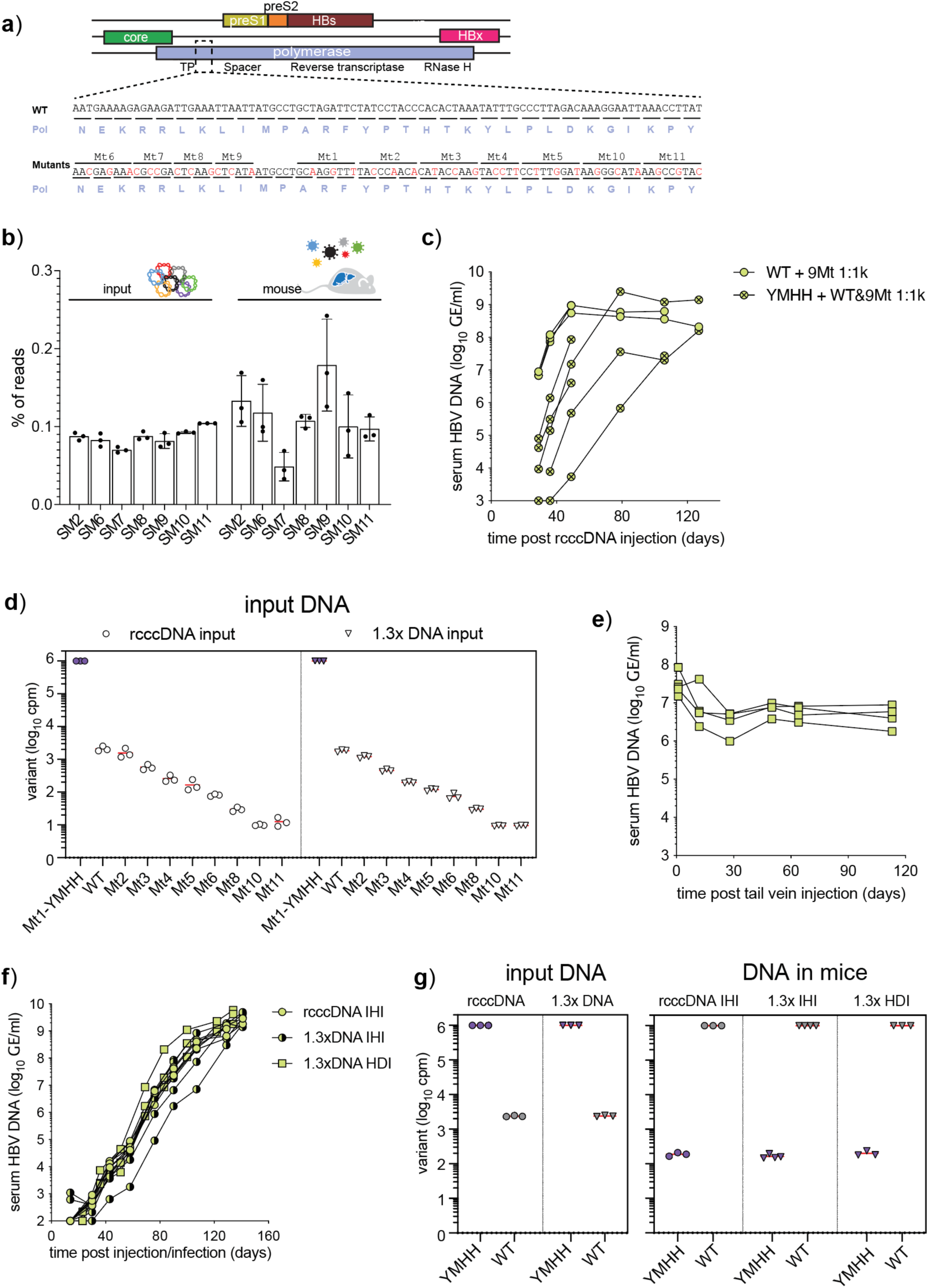
Comparing DNA launch efficiencies across methods. **a)** HBV genome organization with overlapping open reading frames shown in color with nucleotide changes for Mutants (Mt) 1–11 highlighted in red. **b)** An additional 6 mutants (Mt6-11) in the TP domain were spiked into HBV-A_A WT rcccDNA at 1:1,000 and launched using rcccDNA IHI, using silent Mt2 as a positive control. At peak viremia, all mutants, except Mt7 and Mt9, were present at approximately the same frequency as Mt2 and deemed phenotypically silent (input symbols represent technical triplicates, mouse symbols represent individual mice). **c)** huFNRG mice underwent IHI with HBV-A_A WT rcccDNA or YMHH carrier rcccDNA with WT rcccDNA spiked in at 1:1,000. YMHH carrier rcccDNA resulted in delayed viremia kinetics compared to WT rcccDNA (symbols represent individual mice). **d)** Amplicon sequencing of rcccDNA and 1.3× input DNA with the silent mutants spiked into YMHH rcccDNA carrier DNA starting at 1:1,000 at 2-fold dilutions showed the expected dilutions in both input samples (cpm = counts per million, symbols represent technical triplicates). **e)** NRG mice underwent HDI with WT and 8 silent mutants diluted into catalytically dead polymerase (YMHH)1.3x HBV plasmid “carrier” DNA starting at 1:1000 and then 2-fold down (symbols represent individual mice). **f)** huFNRG mice were infected with silent mutant NRG serum or underwent IHI with WT and 8 silent mutant rcccDNA spiked into YMHH carrier DNA, starting with WT at 1:1000 and silent mutants at 2-fold serial dilution. HBV DNA in mouse serum was quantified over time (symbols represent individual mice). **g)** YMDD (WT) region amplicon sequencing of input DNA and mice at peak viremia (cpm = counts per million, input symbols represent technical triplicates, mouse symbols are individual mice).

We designed six additional synonymous mutants in the TP domain (**Sup Fig S3a**), tested their activity as before (**Sup Fig S3b**), and combined the phenotypically silent variants with the previously validated Mt2–5, yielding a panel of eight silent barcodes.

Despite HBV-A_A viremia reaching high levels in huFNRG mice (∼10⁹ GE/ml), reliable detection of rare barcodes is constrained by both limited serum volume and required sequencing depth. To overcome these limitations, we tested whether co-injecting “carrier DNA” with silent mutants at higher dilution ratios could improve sensitivity. YMHH rcccDNA was evaluated as carrier DNA, with WT and the eight mutants spiked in beginning at 1:1,000. At early time points after IHI, mice receiving carrier DNA exhibited viremia that was, on average, approximately 2.5 log_10_ lower than animals injected with WT rcccDNA alone (**Sup Fig S3c**). These results indicated that carrier DNA in the IHI method permits controlled reduction of infectious DNA input, enabling deeper sampling of low-frequency variants.

We then compared all three methods directly (**Fig 3a**). YMHH was linked to Mt1, which was used as carrier DNA, with WT and the eight silent mutants spiked in at two-fold dilutions spanning 1:1,000 to 1:256,000, verified by sequencing the input DNA (**Sup Fig S3d**). The 1.3× DNA containing the eight silent mutants was delivered to NRG mice via HDI, and serum viremia settled in the same range as HDI with undiluted WT DNA (∼5×10^6^ - 1×10^7^ GE/ml), suggesting that other variables limited viremia in this model (**Sup Fig S3e**).

Groups of huFNRG mice underwent IHI or were intravenously infected with HDI-derived NRG serum containing 2×10^6^ GE. All three approaches produced similar viremia kinetics (**Sup Fig S3f**). At peak viremia, serum DNA was sequenced, and silent mutants were quantified using counts per million (cpm) to normalize sequencing depth across samples. All methods supported replication of at least 4,000 variants, as evidenced by the evenly clustered counts of Mt3 across all mice. Starting from a 1:8,000 dilution, however, mutant counts became increasingly uneven (**Fig 3b**). Unexpectedly, cpm of the YMHH variant linked to Mt1 decreased by only approximately 2.5 - 3 log_10_ in the serum of all three groups. To determine whether this reflected low-level DNA recombination, polymerase transcomplementation (26), or PCR contamination, we quantified amplicons containing the YMDD (WT) and YMHH sequences. Input DNA confirmed that WT was spiked into YMHH carrier at 1:1,000, while YMHH in mouse serum was detected nearly 4 log units below WT (**Sup Fig S3g**). Because HDI-infected mice had Mt1 levels similar to those of IHI mice and all were approximately one log above the YMHH serum signal, we conclude that recombination occurred at roughly 0.1% in all methods, while PCR contamination accounted for the ∼0.01% YMHH sequence detected in HDI serum recipients.

These data establish that all three methods can consistently launch at least 4,000 HBV-A_A variants in huFNRG mice, with rcccDNA IHI being the most efficient, supporting up to 32,000 variants across all animals. Given this high launch efficiency and logistical ease, we employed rcccDNA IHI to attempt infection with a broader panel of HBV isolates. All 14 isolates successfully established infection, with only 4 of 52 mice failing to initiate viremia, 3 of which were attributable to early mortality (**Fig 3f**; **Sup Table 2**). Together, these results demonstrate that rcccDNA IHI is the most efficient method for launching HBV in huFNRG mice and can reliably initiate infection across diverse clinical consensus sequences.

### Identification of HBs neutralizing antibody-resistant HBV variants by deep mutational scanning

The high number of variants launched by rcccDNA IHI presents new opportunities. We hypothesized that, among other things, it could support *in vivo* deep mutational scanning (DMS) to identify escape variants against antiviral compounds. To test this, we aimed to test whether this approach could be used to obtain a comprehensive resistance profile for an anti-HBsAg neutralizing monoclonal antibody. We targeted the antigenic loop (AGL) region of the hepatitis B surface (HBs) antigen, a 72-amino acid domain also known as the major hydrophilic region that contains the ‘a’ determinant (aa 124–147) and is essential for infectivity (**Fig 4a**). This region constitutes the principal epitope for neutralizing antibodies (nAbs) and represents a frequent hotspot for immune escape mutations (27). To comprehensively interrogate this region in the HBV-A_A genotype, we constructed a DMS library of 4,608 variants encompassing all 64 possible codons at each amino acid position within the AGL (**Fig 4b, Sup Fig S4a**).

**Figure 4:**
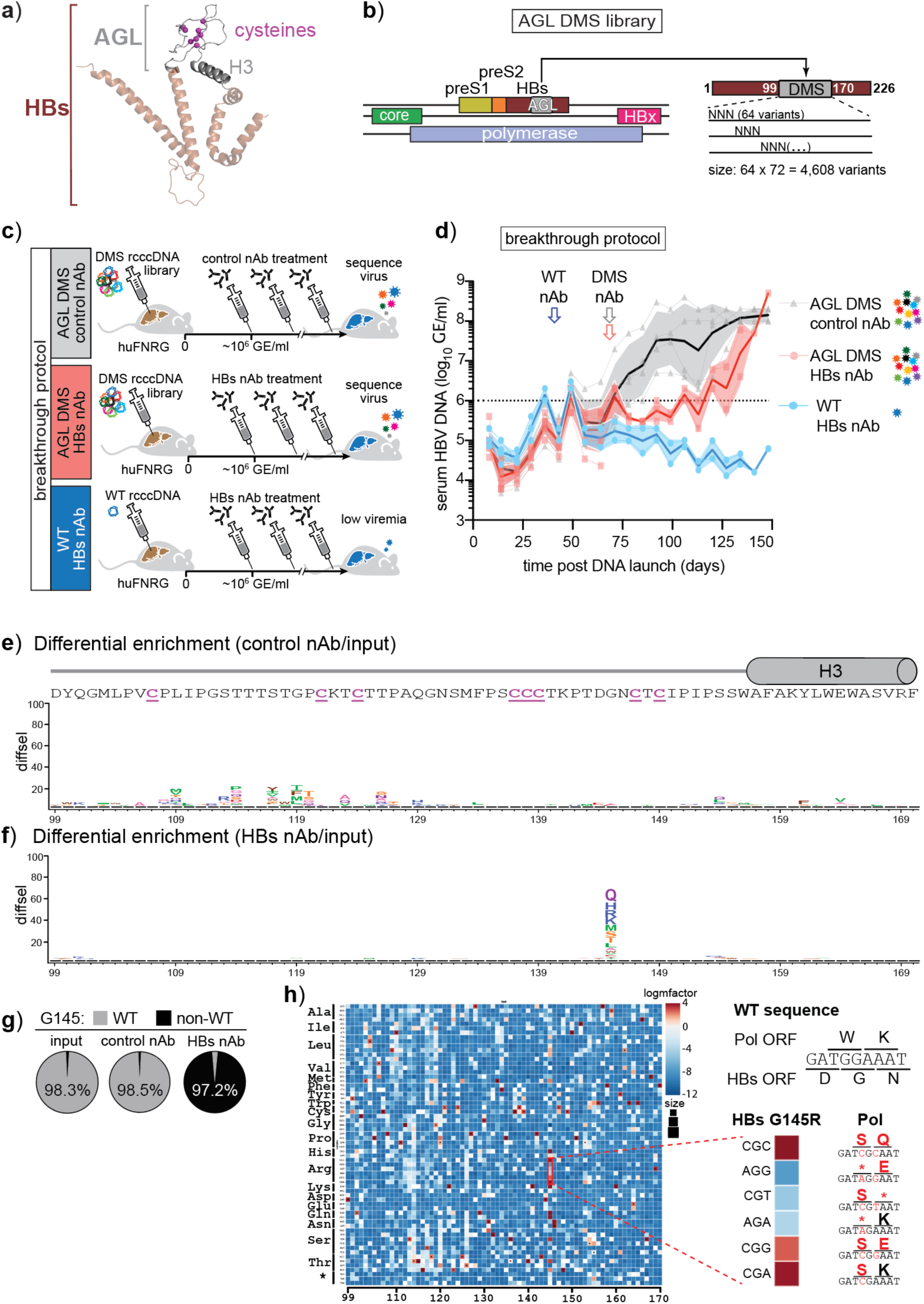
Identification of HBs neutralizing antibody-resistant variants using deep mutational scanning. **a)** Alphafold ribbon structure of HBs monomer with the antigenic loop (AGL) in grey, highlighting the H3 domain and otherwise flexible structure of the AGL containing eight cysteine residues (purple). **b)** Schematic of the antigenic loop deep mutational scanning (DMS) library design. In the HBs open reading frame, each codon from position 99 through 170 was randomized to its 64 possible variants, one codon at a time, for a total of 4,608 variants. **c)** Experimental timeline of the breakthrough protocol. Mice underwent IHI with either WT (blue) or DMS library rcccDNA (grey and red) on day 0 and were monitored for viremia until it reached approximately 1×10^6^ GE/ml. At that time, animals were treated with twice-weekly HBs-neutralizing antibody (nAb) (blue and red) or with a control nAb (grey) until the completion of the experiment. **d)** After IHI, HBV DNA in mouse serum was quantified over time (solid lines means, symbols individual mice, arrows indicate start of nAb treatment for each group). **e)** Amino acid logoplot of differential enrichment in the AGL under control nAb conditions. The wild type amino acid sequence is graphed on top, highlighting the eight cysteine residues (diffsel = log_2_ enrichment relative to the input 1xrcccDNA library). **f)** Amino acid logoplot of enrichments in the AGL under HBs nAb conditions shows multiple amino acid substitutions at G145 were able to break through (diffsel =log_2_ enrichment relative to the input 1xrcccDNA library) **g)** At position G145, the WT sequence dominated in the input DMS library and in the mice under control (ctrl) nAb conditions, while under HBs nAb pressure non-WT sequences dominated. **h)** A log_2_-scaled selection score heatmap of all 64 codon substitutions (Y-axis) across the AGL. Negative (blue) values indicate codon mutants that were not tolerated. Highlighted are the six arginine codons and their impacts on the overlapping polymerase reading frame (black dots indicate WT codon, asterix indicates stop codon).

**Sup Figure S4:**
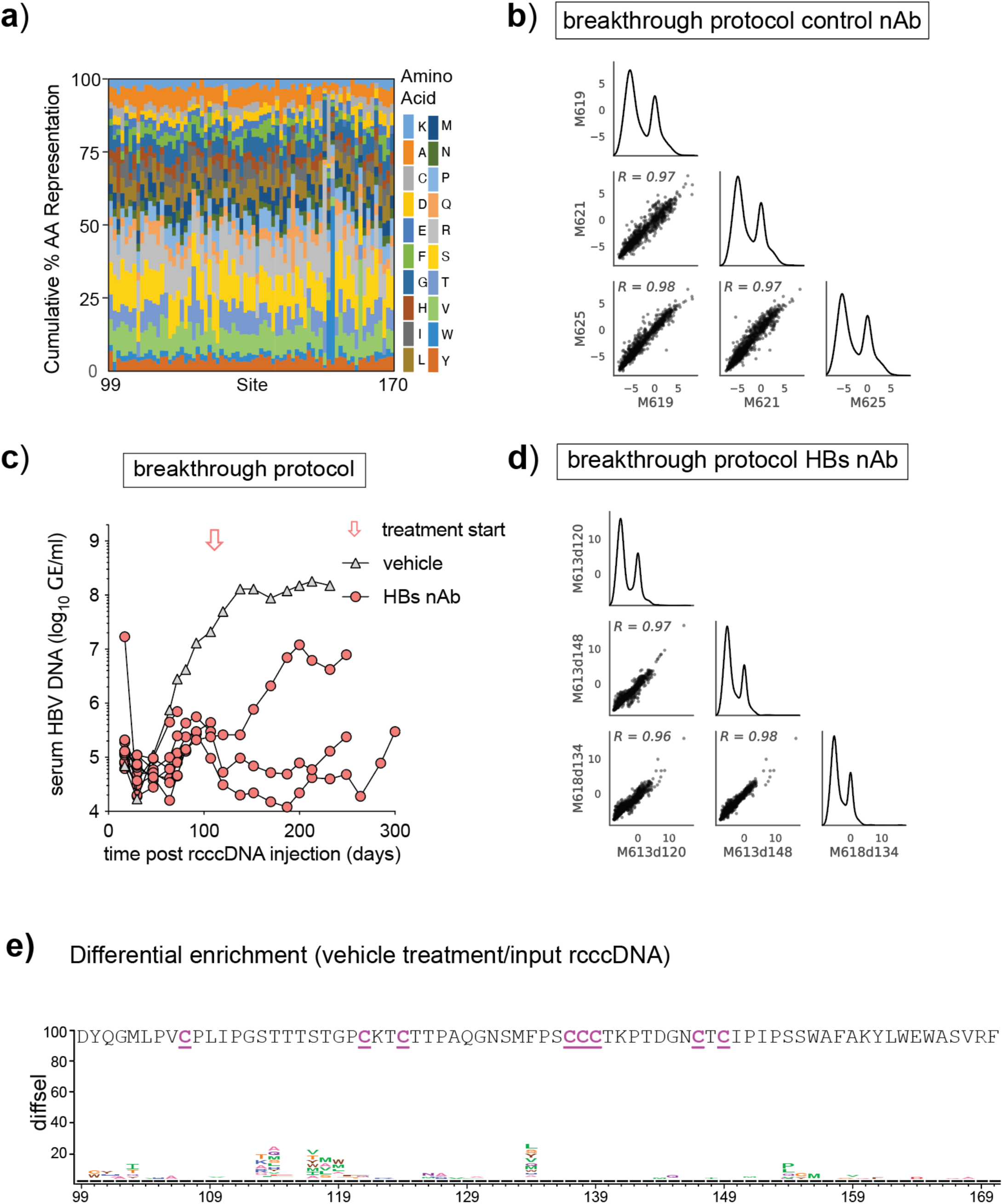
Identification of HBs neutralizing antibody-resistant variants. **a)** Amino acid representation across the codon-randomized library. Stacked bar plot showing the cumulative percentage of non-wildtype amino acid representation at each site. Codon counts were averaged across triplicate input DNA libraries, aggregated by amino acid, and normalized after removing the wild-type amino acid at each position. The resulting plot illustrates the relative composition of variant amino acids across the scanned region. **b)** Reproducibility of control nAb LY1404 differential selection measurements across mice. Selection values were calculated using dms_tools2 and compared across amplicons covering the DMS region, amplified from serum-derived DNA obtained from three individual mice (M619, M621, M625) on day 148 post inoculation with 1xrcccDNA. Each mouse was analyzed with three technical replicates derived from the same amplicon obtained from the input 1xrcccDNA library. Each point represents a single amino-acid mutation. Pearson correlation coefficients are shown for each pairwise comparison. **c)** Repeat breakthrough protocol in huFNRG mice with serum hAlb levels between 4.4 and 15 mg/ml (median 7.7 mg/ml). HBs nAb treatment was initiated on day 111 after IHI, while one mouse near peak viremia received vehicle. Virus in one of the three mice escaped HBs nAb treatment, while the remaining two mice died before reaching viremia levels where variants could be quantified. **d)** Reproducibility of HBs nAb HBc differential selection measurements across samples. Selection values were calculated using dms_tools2 and compared across amplicons covering the DMS region, amplified from serum-derived DNA obtained from three samples from two mice (M613, day 120 and day 148 post inoculation; and M618, day 148 post inoculation with 1xrcccDNA). Each sample was analyzed with three technical replicates derived from the same amplicon obtained from the input 1xrcccDNA library. Each point represents a single amino-acid mutation. Pearson correlation coefficients are shown for each pairwise comparison. **e)** Amino acid logoplot of differential enrichment in the AGL without antibody treatment. The WT amino acid sequence is shown on top, highlighting the eight cysteine residues that form disulfide bonds. The overall pattern of enrichment and regions lacking substitutions is similar to that observed in mice treated with the control nAb, with differences around amino acid 109 (diffsel = log_2_ enrichment relative to the input 1xrcccDNA library).

We first implemented a “breakthrough” protocol in which the complete DMS library was delivered as rcccDNA via IHI, followed by neutralizing antibody pressure to identify variants capable of spread under selection (**Fig 4c**). Once viremia reached approximately 10^6^ GE/mL, we administered HBC24, a nAb targeting the AGL, by intraperitoneal injection (28). Control groups received either LY1404, an irrelevant nAb targeting the SARS-CoV-2 spike protein receptor-binding domain, or were injected with WT HBV-A_A rcccDNA and then received HBC24 treatment at the same viremia threshold. As shown in **Fig 4d**, HBC24 completely blocked viral spread in mice injected with WT virus, whereas the DMS library group showed initial viral suppression followed by breakthrough viremia in HBC24-treated mice, indicating that the AGL variants enabled sampling of sequences that could evade HBC24 neutralizing activity.

To identify the escape variants, we extracted viral DNA from mouse serum and compared variant abundance across treatment groups. The strong correlation in enrichment and depletion patterns between mice treated with the control nAb LY1404 (**Sup Fig S4b**) confirmed consistent library initiation across huFNRG mice. Given the DMS library design, the WT sequence dominated across all randomized regions, indicating no strong selection against it; nevertheless, mild enrichment occurred at multiple positions throughout the AGL, suggesting these variants possess higher relative fitness than WT in the immunodeficient huFNRG environment (**Fig 4e**). A similar enrichment profile in a vehicle-treated mouse from an independent experiment (**Sup Fig S4c,e**) ruled out selection by the control LY1404 nAb as the cause. Conversely, several positions did not tolerate any amino acid substitutions, most notably the eight cysteine residues involved in disulfide bond formation.(29) This demonstrated that IHI effectively delivers a complex DMS library while providing a robust framework to assess the extent of AGL sequence variability permissible under different selective pressures.

HBC24 treatment produced dramatically different results, with strong variant enrichment specifically at the G145 amino acid position, where multiple substitutions disrupted neutralizing activity (**Fig 4f**). G145 glutamine (Q) emerged as the dominant variant, followed by histidine (H) and arginine (R), with strong correlation in variant frequencies between mice (**Sup Fig 4d**). The selection magnitude was striking: non-WT sequences at G145 increased from 1.7% in the input DNA to 97.2% under HBC24 pressure (**Fig 4g**). A global heatmap of codon enrichment confirmed that most substitutions were depleted, while among the six possible G145R codons, only the three that avoided introducing premature stop codons in the overlapping polymerase ORF showed enrichment (**Fig 4h**). This demonstrates that AGL variants were consistently sampled under the dual fitness constraints of overlapping ORFs and HBs nAb pressure, providing a powerful approach for mapping escape mutations in the context of HBV’s genomic architecture.

### Comparing HBs neutralizing antibody-resistant variants between a prophylaxis protocol and the breakthrough protocol

Given the resource-intensive nature of implementing the DMS breakthrough protocol for each experiment or application, we asked whether a simpler, faster prophylactic approach using stocks of diverse viral sera collected from mice could also identify HBs nAb escape variants. We therefore pooled serum from four huFNRG mice infected with the AGL DMS library and treated with control antibody LY1404, and used this as the challenge inoculum. Mice first received two injections of either HBC24 or LY1404, then were intravenously injected with 2×10^6^ GE and monitored for infection (**Fig 5a**). Both groups showed comparable viremia kinetics (**Fig 5b**).

**Figure 5:**
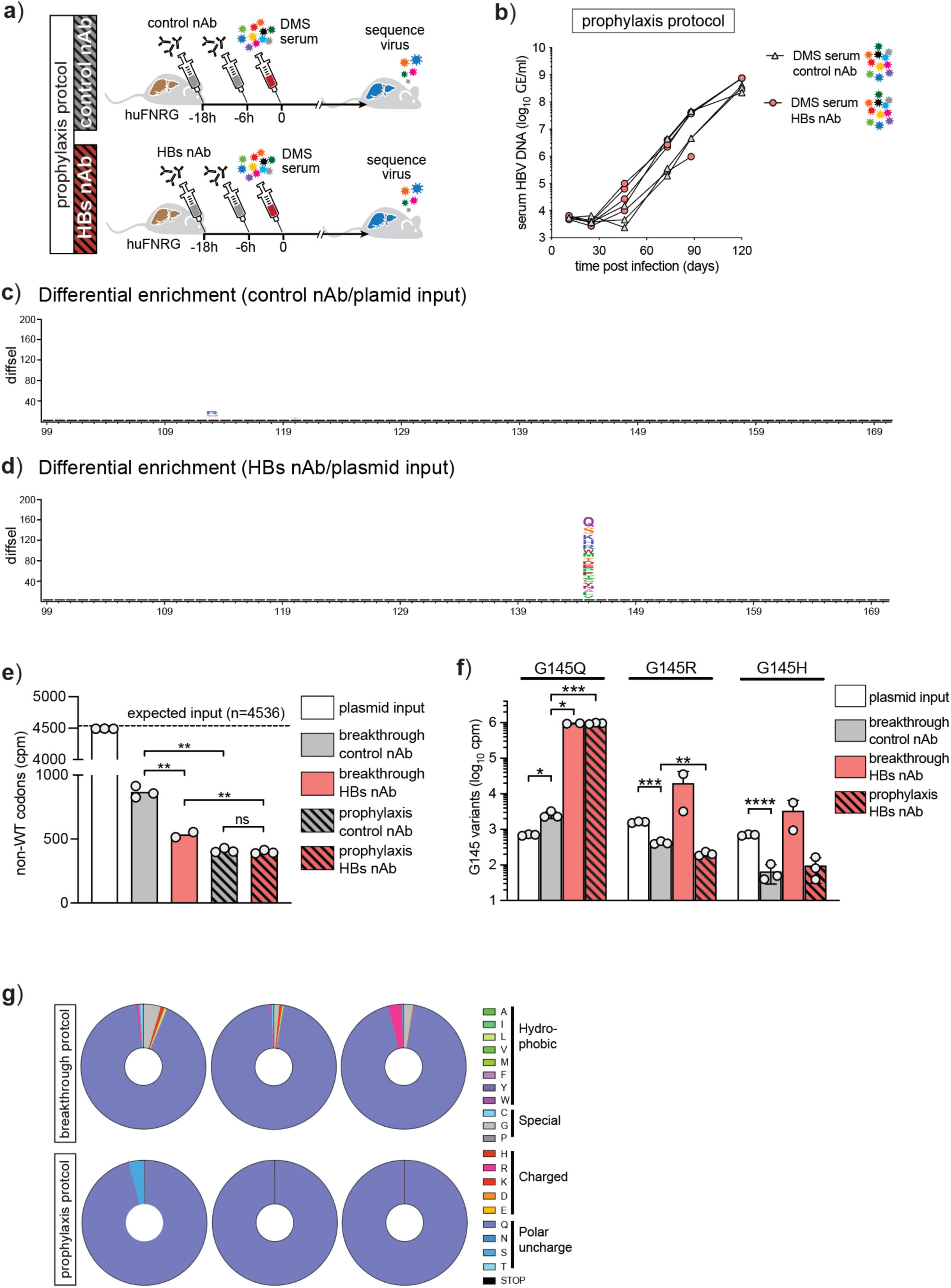
Comparison of HBs-neutralizing antibody-resistant variants between the prophylaxis and breakthrough protocols. **a)** Experimental outline of the prophylaxis protocol. Humanized mice received two injections of control nAb or HBs nAb 18 hours and 6 hours prior to intravenous challenge with pooled DMS serum containing 2×10^6^ GE. Mice were subsequently monitored for viremia. **b)** HBV DNA in mouse serum was quantified over time (symbols: individual mice). **c)** Amino acid logoplot of differential enrichment under control nAb conditions showed that WT residues dominated throughout the AGL (diffsel =log_2_ enrichment relative to the input 1xrcccDNA library). **d)** Amino acid logoplot of enrichments in the AGL under HBs nAb conditions shows that multiple amino acid substitutions at G145 were able to break through (diffsel =log_2_ enrichment relative to the input 1xrcccDNA library). **e)** Comparison of non-WT codons that were detected between protocols and treatments. Input DNA contained 99% of the expected codons, of which, on average, 19% were recovered in the serum from breakthrough mice treated with the control (ctrl) nAb. Under HBs nAb pressure, only 11.9% of non-WT codons were recovered in breakthrough mice. In the prophylaxis protocol, 9.1% and 8.8% non-WT codons were recovered in the control and HBs nAb groups, respectively (codons with >49 read counts were included.For input, symbols represent technical replicates, remaining symbols represent individual mice, ** p<0.01 by Welch t-test, ns = not significant). **f)** Quantification of the three most abundant G145 escape variants in mice with the different treatment protocols (symbols represent input technical triplicates or individual mice), * p<0.05, ** <0.01, *** <0.001, and **** <0.0001 by Welch t-test). **g)** Pie charts showing the distribution of G145 amino acid variants in individual mice from the breakthrough and prophylaxis groups treated with HBs nAb (each pie chart represents one mouse; colors denote amino acids).

DNA extracted from mouse serum at peak viremia was analyzed relative to rcccDNA input and compared across protocols. The prophylaxis protocol with control antibody showed a predominance of WT sequence (Fig 5c), whereas HBC24 treatment again selected G145 variants (Fig 5d). Cross-protocol comparison showed that the breakthrough protocol HBC24 treatment suppressed non-WT variants relative to control, yet both breakthrough conditions contained more variants than prophylaxis mice, consistent with reduced background from non-spreading variants and/or a prophylaxis bottleneck (Fig 5e). Among the three most abundant G145 variants, G145Q was modestly enriched under control antibody treatment and became dominant under HBs nAb treatment in both protocols, whereas G145R and G145H were de-enriched under control conditions and enriched under HBs nAb treatment only in the breakthrough protocol (Fig 5f). These observations suggest that G145Q carries a modest fitness advantage over WT, whereas G145R and G145H carry a fitness cost. All three could nonetheless escape HBs nAb, but the fitter G145Q reached dominance under both protocols, while G145R and G145H did so only in the breakthrough protocol, consistent with established infection better tolerating their fitness cost. WT G145 codons persisted in all breakthrough-protocol mice but disappeared in prophylaxis mice (Fig 5g), indicating that hepatocytes infected before antibody administration continued producing virions throughout the breakthrough experiment.

These results show that the prophylaxis protocol reliably identified dominant HBC24 escape variants but inconsistently captured the full spectrum of escape variants, likely due to bottlenecking or fitness defects in the absence of selective pressure. It did, however, reduce background from variants, most notably WT G145, that expanded before HBC24 selective pressure was applied.

## Discussion

In this study, we found that huFNRG mice were not readily infected with supernatants from hepatoma cell lines or with many clinical samples. Although our observation period may not have been long enough for all isolates, the overall results suggest that most clinical samples and cell line-derived HBV are poorly infectious in this model. Because we detected no viral adaptation in most isolates initiated from DNA in huFNRG mice, we hypothesize that the difficulty in establishing infection arises from inhibitory factors present in some patient plasmas and from the low specific infectivity of virus produced by hepatoma cell lines. This contrasts with an HBV-D strain produced by chimpanzee hepatocytes (30) and HBV-C_Y generated by human hepatocytes in huFNRG mice, both of which exhibited high specific infectivity. Future comparative studies of *in vivo*-derived versus cell culture-derived HBV isolates may help elucidate viral properties that determine infectivity.

The inability to launch a broad range of isolates prompted us to explore alternative approaches, ultimately leading to effective methods for initiating HBV infection directly from DNA. This approach offers several key advantages, some of which were illustrated here. It enables HBV generation from clinical sequences when plasma is unavailable or is poorly infectious. Going forward, any mutation of interest—such as clinically common precore or core promoter mutations—can be evaluated against its isogenic control, enabling more rigorous and controlled investigation of mutation-specific effects. Our protocols also appear more straightforward than a previously reported approach in which an HBV-containing plasmid was first transfected into hepatoma cells to generate stable cell clones, followed by infection of hepatocyte-chimeric mice using supernatants from the highest-producing clones (31). Our most efficient IHI method reliably launched clinical consensus sequences and should facilitate studies of patient-specific HBV isolates.

The use of silent mutants enabled quantification of launch events for the HBV-A_A isolate. One source of uncertainty in the HDI-based estimates is that viremia levels in NRG mice were comparable following injection of undiluted WT 1.3× plasmid DNA and at 1:1,000 dilution. Nevertheless, because silent mutants at the 1:10,000 dilution were detectable in both experiments, we conclude that the HDI approach yields launch estimates comparable to those obtained using the rcccDNA IHI method. We expect that similar results will be true for other isolates, but this remains to be explored.

One promising application of launching large numbers of HBV variants is the ability to conduct *in vivo* DMS experiments. As a proof of principle, we focused on the HBs AGL under nAb pressure. Because all possible codon variants within this region can be represented in a relatively small library (<5,000 variants), this design enabled consistent sampling following rcccDNA IHI. The feasibility of this approach is further supported by our previous demonstration that this mouse model can enrich for HBs nAb escape variants (20). In the presence of an irrelevant nAb, this system provided high-resolution insight into the sequence variation tolerated within the HBs antigenic loop during viral spread in immunodeficient huFNRG mice. Despite long-standing host-virus co-evolution, a substantial number of substitutions were tolerated, although most conferred little apparent fitness advantage over the WT sequence. Notably, substitutions at any of the cysteine residues in the region were absent, consistent with their established role in maintaining AGL structural stability through disulfide bond formation. This includes cysteines 124 and 137, which in some AGL conformations do not form disulfide bonds (29). Under selective pressure from the HBs nAb HBC24, strong and reproducible enrichment was observed at residue G145, a well-established hotspot for antibody escape (32). Multiple amino acid substitutions at this position facilitated viral breakthrough, demonstrating the platform’s capacity to identify escape variants under defined antiviral pressure. Notably, while G145R is the most frequently observed escape mutation in clinical isolates—consistent with its accessibility via a single nucleotide substitution from the WT codon—G145Q requires multiple nucleotide changes and is therefore less readily sampled during natural infection. In contrast, the DMS library equalizes mutational access across all codons. Among enriched variants, only codons compatible with the overlapping polymerase reading frame were selected, highlighting how *in vivo* DMS simultaneously captures constraints imposed by overlapping reading frames and antibody-mediated selection. Together, these findings demonstrate how this approach can resolve context-dependent fitness landscapes shaped by both protein function and genome organization.

Using the DMS library as input, we compared breakthrough and prophylaxis protocols to assess variant tolerance across the viral life cycle. Although most variants were present in the input plasmids, only ∼20% were detected in mice using the breakthrough protocol in the absence of HBC24, underscoring the stringent constraints imposed during viral spread. Even stronger selection was observed with the prophylaxis protocol, likely reflecting reduced carryover of non-spreading variants, additional rounds of selection that enrich optimal over suboptimal codons, and potential bottlenecks during infection establishment. Despite these differences, the WT sequence remained dominant under both protocols, consistent with evolutionary optimization under constraints imposed by overlapping ORFs. In the presence of HBs nAb, both approaches consistently selected for variants at G145, indicating that either protocol can be used to map variant fitness under antiviral pressure. Together, these comparisons provide practical guidance for selecting *in vivo* DMS strategies based on the desired balance between speed and mapping all tolerated variants (breakthrough protocol) versus those that are also fittest in the absence of exogenous selective pressure (prophylaxis protocol).

In summary, we have established robust methods to initiate HBV infection in hepatocyte chimeric mice directly from DNA. The highly efficient IHI approach enables controlled interrogation of viral diversity, including systematic evaluation of mutations, mapping of coding constraints imposed by overlapping reading frames, and analysis of variant fitness under defined selective pressures. This platform should facilitate future studies linking viral sequence variation to replication, immune escape, and therapeutic response, and holds immediate potential for investigating viral variants associated with patient-specific clinical observations and responses to therapeutics.

## Abbreviations

AGL: antigenic loop
cccDNA: covalently closed circular DNA
DMS: deep mutational scanning
GE: genome equivalent
HBs: HBV surface antigen
HBV: hepatitis B virus
HDI: hydrodynamic tail vein injection
huFNRG: human hepatocyte chimeric *Fah^−/−^* NOD *Rag1^−/−^ Il2rg^null^* mice
IHI: intrahepatic injection
IRT: infection and retransplantation
nAb: neutralizing antibody
NRG: immunodeficient NOD *Rag1^−/−^ Il2rg^null^* mice
TRT: transfection and retransplantation
YMHD/YMHH: polymerase dead HBV variants

## Artificial Intelligence use

LLMs (Claude, ChatGPT) were used to modify the original draft, which was written by the authors.

## Author contributions

Y.Y., E.M., L.L.S., F.A.L., C.M.R., W.M.S. and Y.P.J. conceptualized the study. Y.Y., E.M., L.L.S., M.Z., C.Z., C.A.F, G.D., A.A., A.P., A.H., E.R., X.H., C.Q., Y.Z., and Y.P.J. performed the experiments. Y.Y., C.A.F., Y-P.C. performed the analysis. F.A.L., C.M.R., W.M.S and Y.P.J. supervised the research. Y.Y., W.M.S. and Y.P.J. wrote the original draft of the manuscript. All authors reviewed and edited the draft.

## Methods

### HBV isolates

HepDE19 and HepAD38 cells were kindly provided by Dr Haitao Gu (U Pittsburgh, USA) and grown as described.(7, 8) The HBV-A_A plasmid was kindly provided by Drs Amir Shlomai and Yossi Shaul (Weizman Institute, Israel).(22) HBV-C_Y originated from an eAg^−^ patient, injected intravenously into hepatocyte chimeric mice whose serum was stored for subsequent infection experiments.(18) Four de-identified plasma samples from eAg^+^ blood donors that contained >1.7×10^8^ IU/ml HBV DNA were obtained from the New York Blood Center.(33) DNA of 25 clinical consensus sequences(24) across genotypes A through F was synthesized by Twist Bioscience. HBV DNA was quantified by Taqman as described(34) using genotype compatible primers/probes.

### Mice

*Fah^−/−^* NOD *Rag1^−/−^ Il2rg^null^*(FNRG) mice were created by backcrossing *Fah^−/−^* liver injury mice(35) to NOD *Rag1^−/−^ Il2rg^null^* (NRG, Jackson Laboratories) animals for 13 generations.(17) To create humanized (hu)FNRG mice, animals were preconditioned with retrorsine and withdrawn from nitisinone (Yecuris) prior to transplantation with 5×10^5^ primary human hepatocytes from donor HUM4188 (Lonza).(21) Animals were cycled on nitisinone for 10-12 weeks after which human albumin (hAlb) was quantified in mouse serum using a species-specific ELISA (Bethyl Labs). Animals with hAlb levels >9mg/ml were used for intrahepatic injection, and animals with hAlb levels >3mg/ml for infections. All procedures were approved by the Rockefeller University IACUC under protocols 18063, 21056 and 24022.

### Generation of HBV culture supernatants

HBV was produced using HepAD38 and HepDE19 cells as described.(33) Briefly, cells were cultured in DMEM supplemented with 10% fetal bovine serum (FBS), 1% non-essential amino acids (NEAA), and 1 μg/mL tetracycline prior to confluency. Upon reaching confluency, cells were switched to DMEM containing 3% FBS and 1% NEAA without tetracycline (−Tet) to induce HBV replication. Culture supernatants were collected every 2–3 days for up to four weeks, and virus-containing media were concentrated using Centricon Plus-70 centrifugal filter devices (Cat. No. UFC710008; MilliporeSigma). Concentrated viral stocks were aliquoted and stored at −80 °C until use. Based on temperature dependent maturation to facilitate heparan sulfate binding,(36) HepDE19 supernatants were harvested and incubated at 37°C for 20hrs (warm), frozen at −80°C (cold) or combinations prior to injection into huFNRG mice.

### Generation of mutants and conversion to rcccDNA

All HBV genotype plasmid sequences used in this study have been deposited in GenBank (Table S2). The mt1–mt11 constructs each harbor three silent nucleotide substitutions in the TP domain, introduced into a wild-type 1.3× HBV-A_A.(22) YMHD and YMHH variants were generated by mutating the YMDD catalytic motif within the reverse transcriptase subdomain of the polymerase in the same plasmid backbone. Genotypes B, C, D, and E-H were described and into the pGEM-3Z - backbone. AGL-DMS library were synthesized that has the saturated mutations at the 99-170 amino acid at the HBs region.

### Converting 1.3X plasmid into rcccDNA

The 1.3× HBV plasmids were prepared using an EndoFree Plasmid Maxi Kit (Qiagen, Cat#12362). To generate at least 10 µg of rcccDNA, 100 µg of plasmid DNA was digested in a 500 µl reaction containing 200 U PciI (to cut the HBV repeat region) and 200 U BglI (to cut the plasmid backbone). Digestion products were separated on a 1% TAE agarose gel, and the ∼3.2 kb band was excised and purified. Ligation was performed at a DNA concentration of 2 ng/µl in 1× ligation buffer using 0.5–1 U/µl DNA ligase overnight.

Typically, large-scale ligation reactions were performed in 15 ml total volume with 30 µg linear DNA and 15,000–30,000 U ligase at room temperature overnight. On day 2, 5 volumes of PB buffer (Qiagen) were added, and DNA was purified using QIAquick Gel Extraction columns (up to 10 µg DNA per column).

Purified DNA was analyzed on a 1% TAE agarose gel, and the ∼2.1 kb band corresponding to 1× rcccDNA was excised and purified using the same kit. DNA was eluted in 30 µl ddH₂O per column and diluted to a final concentration of 15 ng/µl in 200 µl 1× PBS.

### Retransplantation launch

Mouse-passaged primary human hepatocytes (mpPHH) were isolated from huFNRG mice and plated on 6 well collagen coated plates (BD Bioscience) as described.(21) One day after plating, media was changed to hepatocyte complete medium (HCM, Lonza). For infection and retransplantation, mpPHH cultures were infected with HepDE19 supernatants or HBV-C_Y serum. For transfection and retransplantation, mpPHH cultures were transfected with 1.5 µg of rcccDNA per well using lipo2000 transfection reagent. 2 days later, mpPHH were mobilized with TripleE express and 3×10^5^ cells were transplanted into huFNRG mice.

Intrasplenic transplantation was performed as described.(17) For intrahepatic transplantation, huFNRG mice were anesthetized with isoflurine, abdominal skin was shaved and sterilized with iodine followed by 70% isopropanol. An approximately 1cm subcostal incision was made and the large lobe was mobilized onto the abdominal wall. Using a 31G insulin syringe (BD Bioscience), mpPHH resuspended in 70μl of PBS were injected in 5-7 locations of the liver. Pressure was applied with sterile cotton tips after each injection to prevent bleeding. After the last intrahepatic injection, the peritoneum was closed using Vicryl sutures, the skin using metal clips and bupivacaine was applied topically. Animals received buprenorphine 0.05mg/kg for postsurgical analgesia.

### Intrahepatic DNA injection launch

For intrahepatic injection, highly humanized huFNRG mice were anesthetized with isoflurine, abdominal skin was shaved and sterilized with iodine followed by 70% isopropanol. An approximately 1cm subcostal incision was made and the large lobe was mobilized onto the abdominal wall. Using a 34G syringe (Hamilton company 7633-01& 207434-10), DNA resuspended in 100μl of PBS was injected in ∼20 locations throughout the large lobe. Pressure was applied with sterile cotton tips after each injection to prevent bleeding. After the last intrahepatic injection, the peritoneum was closed using Vicryl sutures, the skin using metal clips and bupivacaine was applied topically. Animals received buprenorphine 0.05mg/kg for postsurgical analgesia.

### Hydrodynamic tail vein injection launch

NRG mice were weighed and placed in a constrainer. The tail vein was warmed under a heatlamp and 20μg of DNA was diluted in PBS to total volume that corresponded with 9% body weight. Using a 3ml syringe (BD Bioscience) and 27G needle, the tail vein was cannulated and the total volume injected over 3-5 seconds. Mice were allowed to recover in cage on a heatpad until they started ambulating.

### Neutralizing antibody administration

For the breakthrough protocol, animals that had undergone rcccDNA intrahepatic injection were bled weekly. When a group of mice reached approximately 1×10^6^ GE/ml, murinized HBC24 or LY1404 were resuspended at 1mg/ml in PBS and 0.5ml was injected intraperitoneally twice weekly for the duration of the experiment. For the prophylaxis protocol, huFNRG mice received two injections of 0.5mg 18 and 6 hours prior to intravenous challenge with mouse serum containing 2×10^6^ GE HBV. They were monitored for viremia at the timepoints indicated.

### Sequencing and analysis

40 µl of the serum was used for DNA extraction using the DNA Blood Kit (Qiagen, 51105). PCR0 was performed to amplify a near full-length HBV template as described^12^ using KOD Xtreme Hot Start DNA Polymerase (Sigma, 71975-3). The number of PCR cycles was determined based on the HBV DNA copy number quantified by the Taqman assay described above using the equation (y=−1.864×ln(x)+45.336), where y is the number of PCR cycles, and x is the HBV copy number in 5 µl DNA sample. The equation was derived from previous experience and further refined by empirical optimization based on the yield of PCR products visualized by SYBR Safe staining of agarose gels. At least 10^6^ genome equivalents (GE) of HBV DNA were used for DMS library construction and multiple PCR0 were combined to obtain sufficient template for library construction depending on HBV DNA levels in the serum. PCR1 was performed using RU-O-33251 (CTTTCCCTACACGACGCTCTTCCGATCTNNNNNNNNTATCGCTGGATGTGTCTGCGG) as the forward primer and RU-O-33242 (GGAGTTCAGACGTGTGCTCTTCCGATCTNNNNNNNNCAAATGGCACTAGTAAACTGAGCCAAGA) as the reverse primer to generate an amplicon containing the HBs antigenic loop (AGL). PCR2 was performed under bottleneck conditions with 350,000 input templates, as previously described. Pooled libraries were sequenced at the Rockefeller University Genomics Resource Center using either the Illumina or AVITI platform with paired-end 250bp sequencing (PE250). Fastq files were analyzed using dms2_tools2, developed by the Bloom laboratory (https://github.com/jbloomlab/dms_tools2). Correlation plots were generated using dms2_batch_diffsel, using the arguments that summarizes the positive differential selection (positive mutdiffsel) across samples. Sequence logo plots were generated using dms2_logoplot. Heatmaps were generated from codon tables produced by dms_tools2 using the analysis pipeline available at https://github.com/HBV-DMS/R1/blob/main/JNote.ipynb.

---

**Supplemental Table 1:**
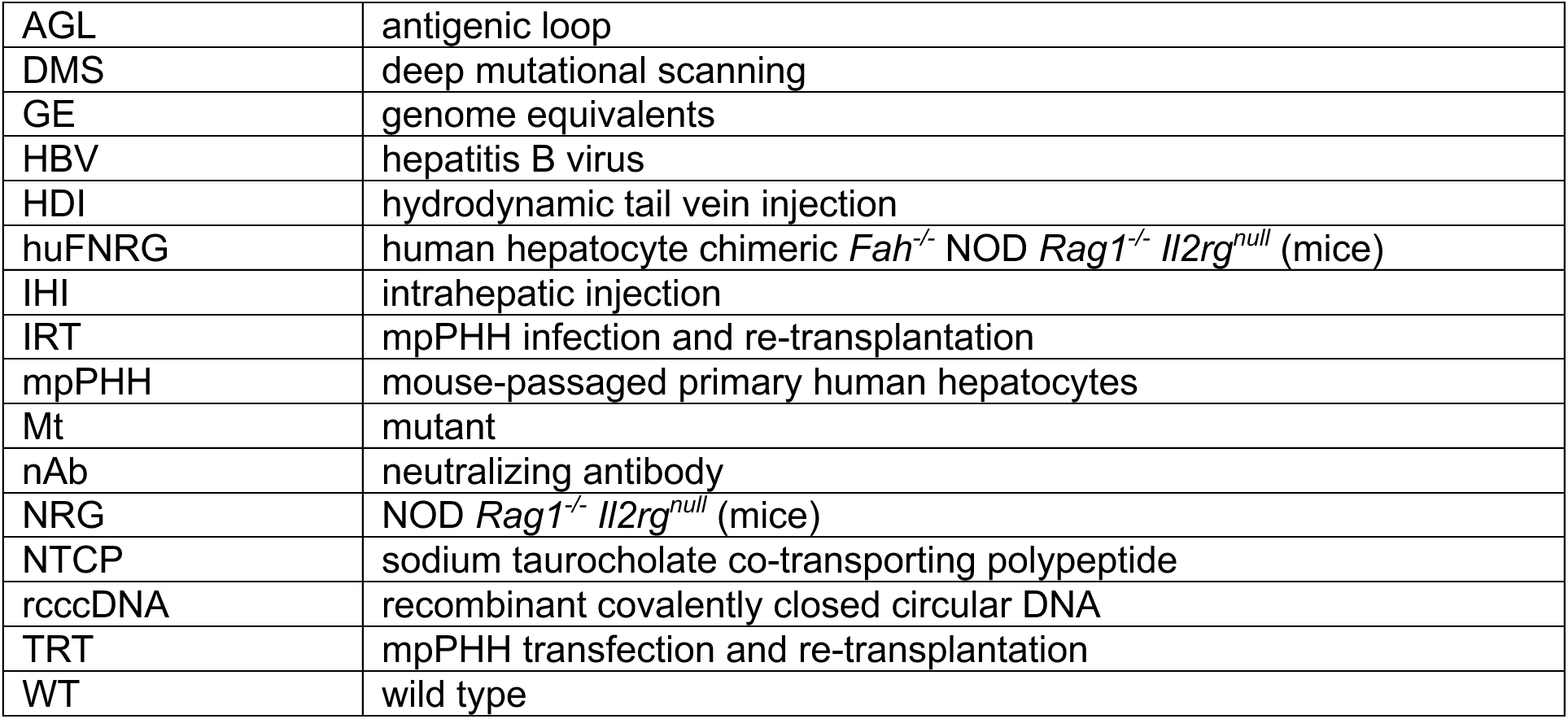
Abbreviations.

**Suplemental Table 2:**
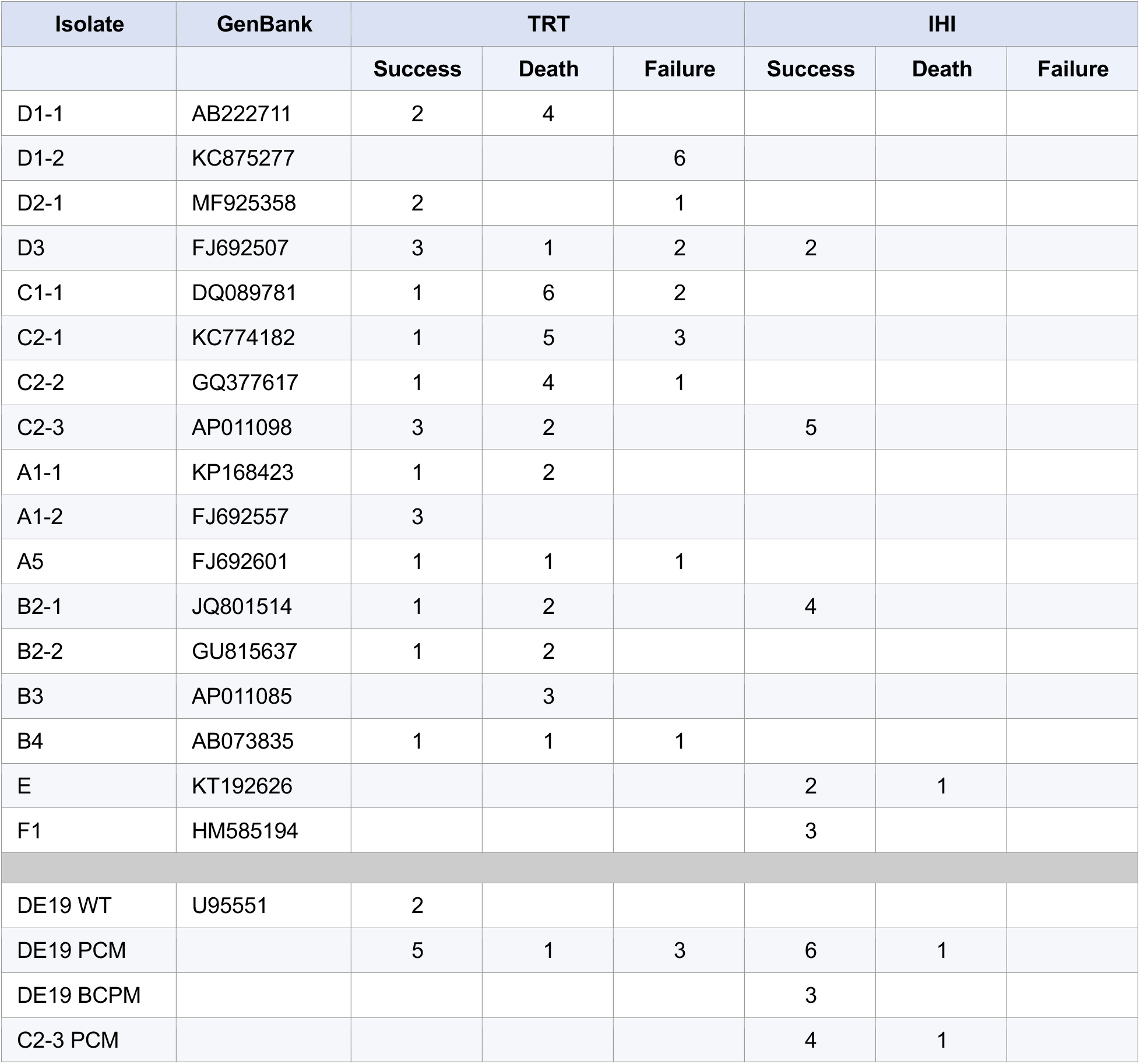
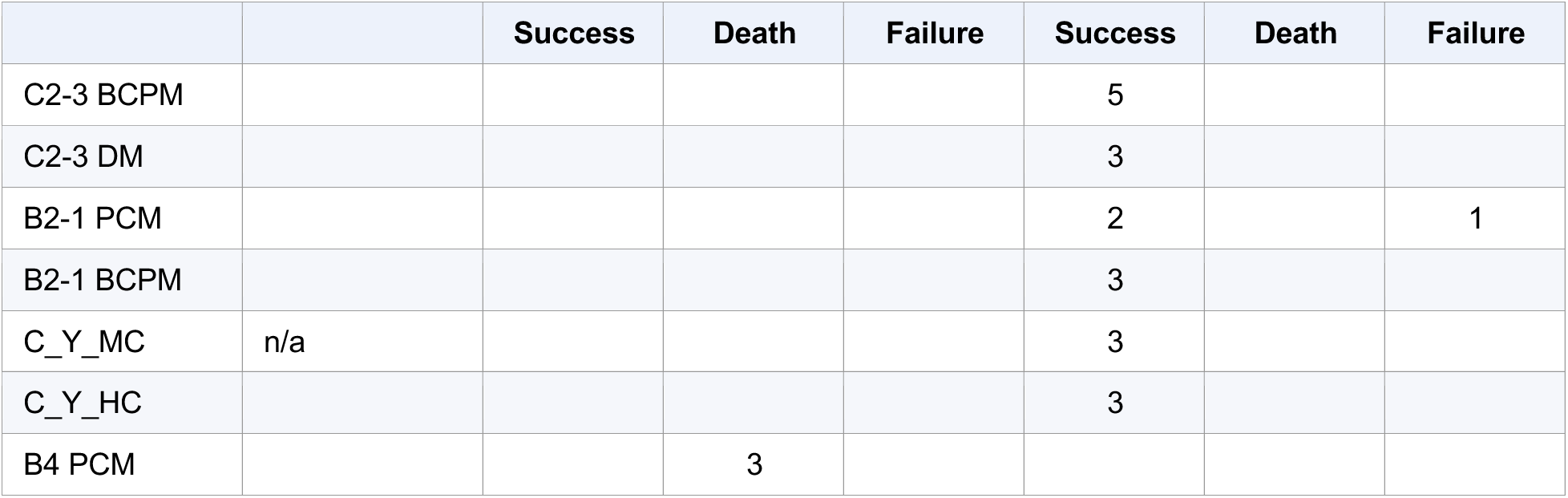
HBV Isolate Infection Launch Summary.

